# Age- and Sex- Driven Transcriptional and Metabolic Diversity in Myeloid-Derived Suppressor Cells After Mouse Sepsis

**DOI:** 10.1101/2025.10.06.680736

**Authors:** Christine Rodhouse, Dayuan Wang, Hongru Tang, Valerie E. Polcz, Miguel Hernández-Ríos, Xuanxuan Yu, Leandro Balzano-Nogueira, Jaimar C. Rincon, Angel Charles, Seiichiro Fujishima, Quan Vo, Marvin L. Dirain, Ricardo Ungaro, Dina C. Nacionales, Leilani Zeumer-Spataro, Larissa Langhi Prata, Shawn D. Larson, Gemma Casadesus, Feifei Xiao, Shannon M. Wallet, Maigan A. Brusko, Tyler J. Loftus, Letitia Bible, Alicia M. Mohr, Paramita Chakrabarty, Michael P. Kladde, Clayton E. Mathews, Lyle L. Moldawer, Robert Maile, Guoshuai Cai, Philip A. Efron

## Abstract

Sepsis induces profound immune dysregulation, often resulting in chronic critical illness characterized by persistent immunosuppression and poor outcomes. Myeloid-derived suppressor cells (MDSCs) are central mediators of this immunosuppressive phenotype, yet the influence of age and sex on their transcriptional and metabolic states remain poorly understood. Here, we employed single-cell RNA sequencing of splenic leukocytes from young (3-4 months) and older (18-24 months) adult male and female mice subjected to a clinically relevant murine sepsis model to define age- and sex-specific MDSC phenotypes. We identified significant differences regarding age and sex in MDSC expansion, transcriptome, canonical pathway activation, RNA velocity, mitochondrial metabolism, and predicted cell-cell communication after sepsis. Using drug2cell analysis of total leukocytes we also identified cohort-specific drug target profiles. These findings underscore the importance of age and sex in shaping sepsis-induced MDSC biology and suggest that personalized immunomodulatory strategies targeting MDSCs could improve sepsis outcomes.

## Introduction

Sepsis, defined as organ dysfunction/failure in response to an infection^1^, is a disease that remains costly, morbid, and frequently fatal^1^. Although acute in-hospital mortality has decreased due to earlier diagnosis and optimized clinical therapy (e.g. Surviving Sepsis Campaign)^2^, long-term outcomes remain unacceptably poor^3,4^. Among septic patients who survive the initial *“cytokine storm”* and avoid early organ injury and death, two predominant clinical trajectories have emerged: 1) those who are successfully managed, rapidly recover, and are discharged from the ICU and, 2) those who remain critically ill with sustained organ dysfunction, developing chronic critical illness (CCI), leading to either death or persistent physical and cognitive impairment^3,5,6^. Sepsis survivors who develop CCI are more likely to experience infections and sepsis recidivism, disability, and death in the first year^3,5–8^. What drives this failure to fully recover remains unclear, but CCI patients often develop an immunological endotype described as the Persistent Inflammation, Immunosuppression and Catabolism Syndrome (PICS)^7^.

Myeloid-derived suppressor cells (MDSCs) expand in number and are associated with poor outcomes in sepsis survivors^9,10^. First described in the cancer literature in the early 2000s^11^, MDSCs arise during pathological states of chronic immune activation in the host, and contribute to chronic inflammation while simultaneously suppressing leukocyte antimicrobial functions, particularly in lymphocytes^12^. MDSCs can be phenotypically classified in three subsets: granulocytic/polymorphonuclear (PMN)-MDSCs, monocytic (M)-MDSCs, and early (E)-MDSCs, each exhibiting unique immunosuppressive mechanisms^13^.

One of the major barriers in the clinical translation of immunomodulatory therapies for sepsis has been the failure to incorporate principles of personalized medicine^14^. A growing body of evidence indicates that host-specific factors, particularly age and sex, significantly influence the immunological response to sepsis and the likelihood of recovery^15–19^. For example, although some controversy remains, young females generally exhibit more favorable outcomes following sepsis when compared to age-matched males or older adults of either sex^15–19^.

Sex- and age-dependent disparities in sepsis outcomes likely reflect fundamental differences in immune activation, resolution, epigenetics, mitochondrial metabolism and repair, and cell differentiation^20–25^. With respect to MDSCs, both age and sex are known to modulate the hematopoietic landscape and the myeloid compartment. Aging, in particular, is associated with myeloid skewing, impaired resolution of inflammation, and accumulation of dysfunctional myeloid cells^19,26–29^. These differences underscore the importance of stratifying patients not only by disease severity but also by age and sex to optimize immunotherapeutic strategies in sepsis. Given that the dysregulated expansion, activation, and persistence of MDSCs have emerged as a potential therapeutic target in sepsis^30^, it is essential to delineate how pathological immune responses vary across different cohorts. Defining these context-specific immune trajectories will be critical for establishing individualized baselines of immune homeostasis and for designing tailored immunotherapies that account for age- and sex-related differences.

To investigate the mechanistic basis underlying age- and sex-related disparities in sepsis outcomes, we employed a validated murine model of surgical sepsis that reliably induces PICS^19,31^ and more closely recapitulates the septic human condition^31^. Mice were selected as a preclinical model due to their well-characterized immune systems, the feasibility of acquiring age- and sex-matched cohorts, and the ability to control for confounding variables that are challenging to address in human studies: namely, pre-existing conditions, timing of sepsis onset, and access to tissue-level single-cell data. This study aimed to define, at single-cell resolution, the transcriptomic and metabolic features of splenic MDSCs from young and older adult male and female mice following sepsis. Additionally, mitochondrial activity was analyzed due to its established links to cell differentiation, proliferation, survival, epigenetics regulation, and immune cell function^32–36^ We hypothesized that age and sex influence the expansion, phenotype, and function of MDSCs during sepsis. This approach also enabled precise dissection of how age and sex shape MDSC biology in sepsis and can support the development of more targeted immunomodulatory therapies.

## Results

### 3.1 Sepsis vs naïve

### 3.1.1 Expansion of MDSCs following sepsis

We performed scRNAseq of splenic myeloid cells across age-, sex-, and disease-stratified murine cohorts. For our initial analysis we used canonical gene markers^37^ to identify transcriptionally distinct subsets of MDSCs, including M-MDSC, PMN-MDSC, and E-MDSC populations, as visualized in the UMAP and dot plots shown in **Fig. S1**. Specifically, **Fig. S1A** and **S1C** display the low-dimensional projection of annotated myeloid populations, while **Fig. S1B** and **S1D** highlight canonical and dataset-specific marker genes that validate the identity of each MDSC subtype. Quantitative analysis demonstrated a significant expansion of all three MDSC subsets in septic mice compared to naïve controls, with differential distributions across cohorts (**Table S2, Table S3, Fig. 1A**).

**Figure 1.**
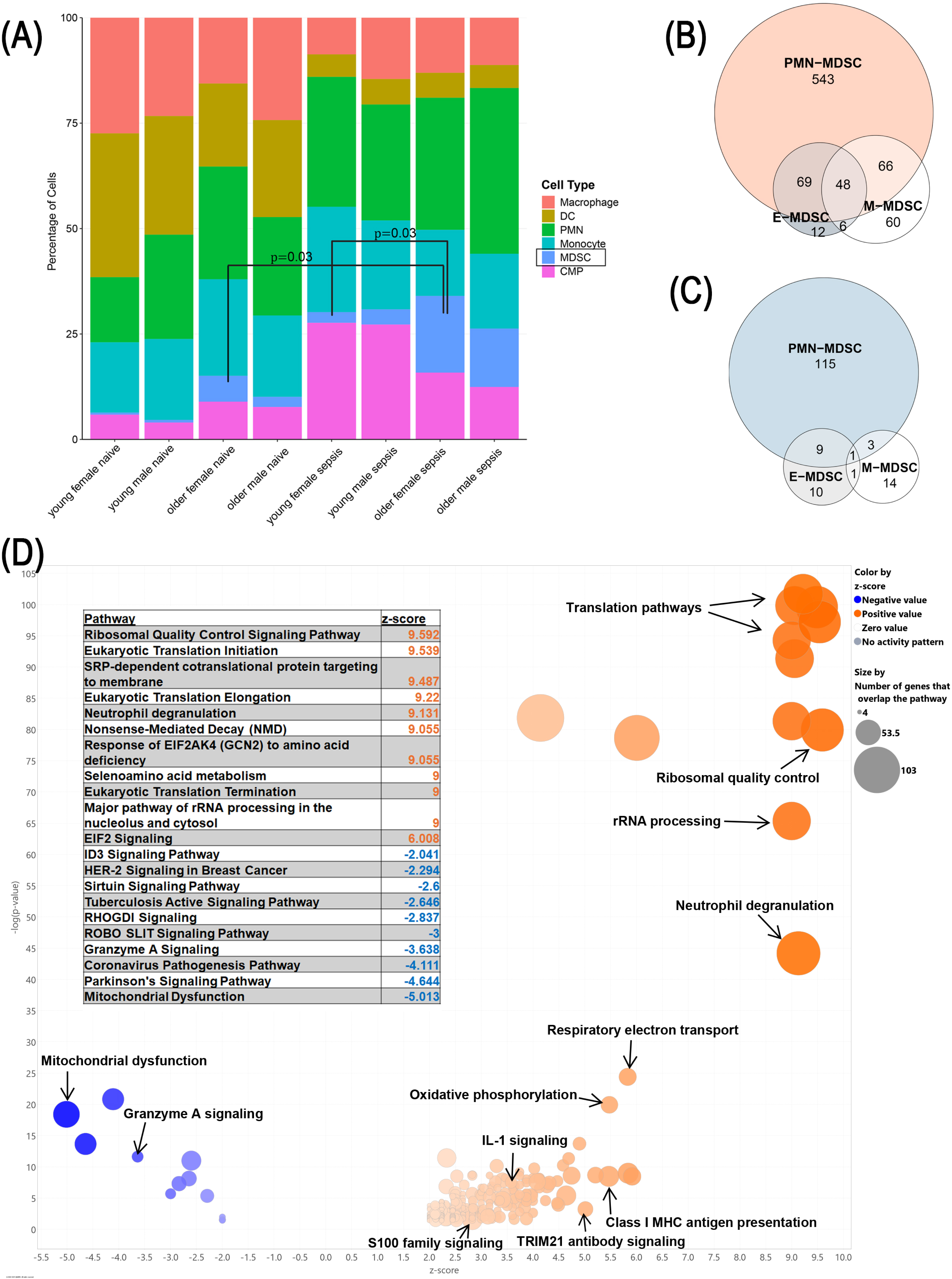
Cell proportion and transcriptomic analysis of MDSCs in septic vs naïve mice. **(A)** There was a significant expansion of MDSCs in all septic mice compared to naïve. Additionally, there was a greater expansion of MDSCs in the septic older mice compared to young but no significant difference between the sexes. **(B/C)** A differential expression analysis (DE) was performed between all the septic mice and all the naïve mice. Panel **B** shows the upregulated DEG overlap between the MDSC subtypes and panel **C** the downregulated gene overlap in sepsis compared to naïve. **(D)** Pathway analysis was performed using IPA. There was mostly upregulation (orange) of the canonical pathways in sepsis compared to naïve. IPA analysis of the PMN-MDSCs for this comparison is shown to represent all the MDSCs. There was upregulation of pathways related to translation and ribosome function, immune regulation like neutrophil degranulation and class I MHC antigen presentation, as well as those related to mitochondria function like oxidative phosphorylation and electron transport. The most significant downregulated pathways included those related to mitochondrial reactive oxygen mediated stress and mitochondrial death (granzyme A). Dendritic cell (DC), common myeloid progenitor (CMP).

#### 3.1.2 Transcriptomic and canonical pathway reprogramming of MDSCs following sepsis

For our initial analysis, we compared pooled sepsis and naïve cohorts, which included animals of both sexes and across age groups, to establish a global overview of how sepsis alters MDSC transcription and canonical pathway activation. This approach allowed us to identify the dominant transcriptomic and functional shifts associated with sepsis independent of host-specific variables, serving as a foundational reference point before dissecting age- and sex-specific effects in subsequent analyses. We conducted differential gene expression (DEGs) analysis for each MDSC subtype to evaluate transcriptomic alterations between septic and naïve mice. The majority of the DEGs were observed to be significantly (FDR q<0.01) upregulated in septic versus naïve mice, indicating broad activation of transcriptional programs associated with sepsis (**Table 1**, **Fig. 1B/C**). PMN-MDSCs in particular had the most DEGs with 543 upregulated (**Fig. 1B**) and 115 downregulated (**Fig. 1C**). IPA revealed that the most enriched canonical signaling pathways were also upregulated in septic mice, consistent with a pro-inflammatory, immunoregulatory, and stress-adaptive transcriptional shift (**Fig. 1D**). Notably, the most significantly downregulated canonical pathways in septic MDSCs compared to naïve MDSCs were associated with mitochondrial stress responses.

**Table 1.** DEG Analysis. Number of significant DEGs per adult mouse cohort and whether they were up- or downregulated based on the log2FC. YF = young female, YM = young male, OF = older female, OM = older male.

| Comparison | Direction | Number of Differentially Expressed Genes |  |  |
| --- | --- | --- | --- | --- |
|  |  | E-MDSC | M-MDSC | PMN-MDSC |
| Sepsis vs naïve | up regulation | 135 | 180 | 726 |
|  | down regulation | 21 | 19 | 128 |
| Young vs older sepsis | up regulation | 4 | 15 | 48 |
|  | down regulation | 6 | 140 | 29 |
| Female vs male sepsis | up regulation | 2 | 4 | 3 |
|  | down regulation | 1 | 5 | 0 |
| YF vs YM sepsis | up regulation | 2 | 1 | 3 |
|  | down regulation | 1 | 4 | 1 |
| OF vs OM sepsis | up regulation | 1 | 5 | 1 |
|  | down regulation | 5 | 153 | 2 |
| YF vs OF sepsis | up regulation | 1 | 6 | 3 |
|  | down regulation | 2 | 1 | 5 |
| YM vs OM sepsis | up regulation | 0 | 232 | 2 |
|  | down regulation | 3 | 55 | 4 |

### 3.2 Age and sex comparisons within septic hosts

#### 3.2.1 Age-dependent expansion (and sex-associated trends) of MDSC subtypes following sepsis

Next, we quantified the relative abundance of MDSC subtypes across age, sex, and disease groups (**Table S2, Table S3, Fig. 1A**). Under homeostatic conditions, naïve young adult mice exhibited very low numbers of splenic MDSCs, consistent with restrained myelopoiesis in the absence of inflammatory stimuli (**Fig. 1A**). In contrast, naïve older adult mice displayed a significant expansion of MDSCs, driven primarily by PMN-MDSCs, aligning with prior reports (e.g. inflammaging) (**Table S2**)^38^. Although all three MDSC subtypes demonstrated marked expansion in both age groups after CLP+DCS, older adult septic mice harbored proportionally and absolute greater numbers of MDSCs (versus young adult counterparts). However, no significant differences in MDSC proportions were observed between exclusively male and female mice in either the naïve or septic conditions (**Table S3**).

There were also combined age and sex differential expansion of MDSC subtypes after sepsis. Septic young adult female mice exhibited significantly lower frequencies of M- and PMN-MDSCs compared to septic older adult females (*p*<0.05; **Fig. 1A, Table S4**). When compared to age- and sex-matched naïve controls, older adult septic females had significant increases in both M- and PMN-MDSCs (*p*<0.05), while older adult males only exhibited significantly increased M-MDSCs following sepsis compared to their age- and sex-matched naïve controls (*p*=0.01).

#### 3.2.2 Age- and sex-specific differential transcriptomic and canonical pathway reprogramming of MDSCs following sepsis

To investigate how host age and sex influenced MDSC gene expression profiles during sepsis, we performed stratified transcriptomic analyses across cohorts. DEG analysis revealed that M-MDSCs from young adult septic mice had an increased number of downregulated genes. In addition, PMN-MDSCs from young adult septic mice exhibited a greater number of significantly upregulated genes than their older adult counterparts (**Table 1**). In contrast, E-MDSCs demonstrated relatively few age-related DEGs, suggesting a more limited transcriptional responsiveness of this myeloid subset to host age.

IPA further demonstrated distinct age-dependent reprogramming (**Fig. 2A**). Young adult septic animals exhibited significant upregulation of pathways involved in mitochondrial biogenesis and function relative to older septic mice. Conversely, pathways associated with mitochondrial dysfunction and granzyme A signaling were significantly downregulated in young adult septic animals. Transcriptional and translational machinery were also elevated in all MDSC subsets from young adult septic mice relative to older septic mice. Immune and inflammatory signaling pathways displayed a more complex pattern; most were downregulated in young adult animals, including innate-adaptive immune communication.

**Figure 2.**
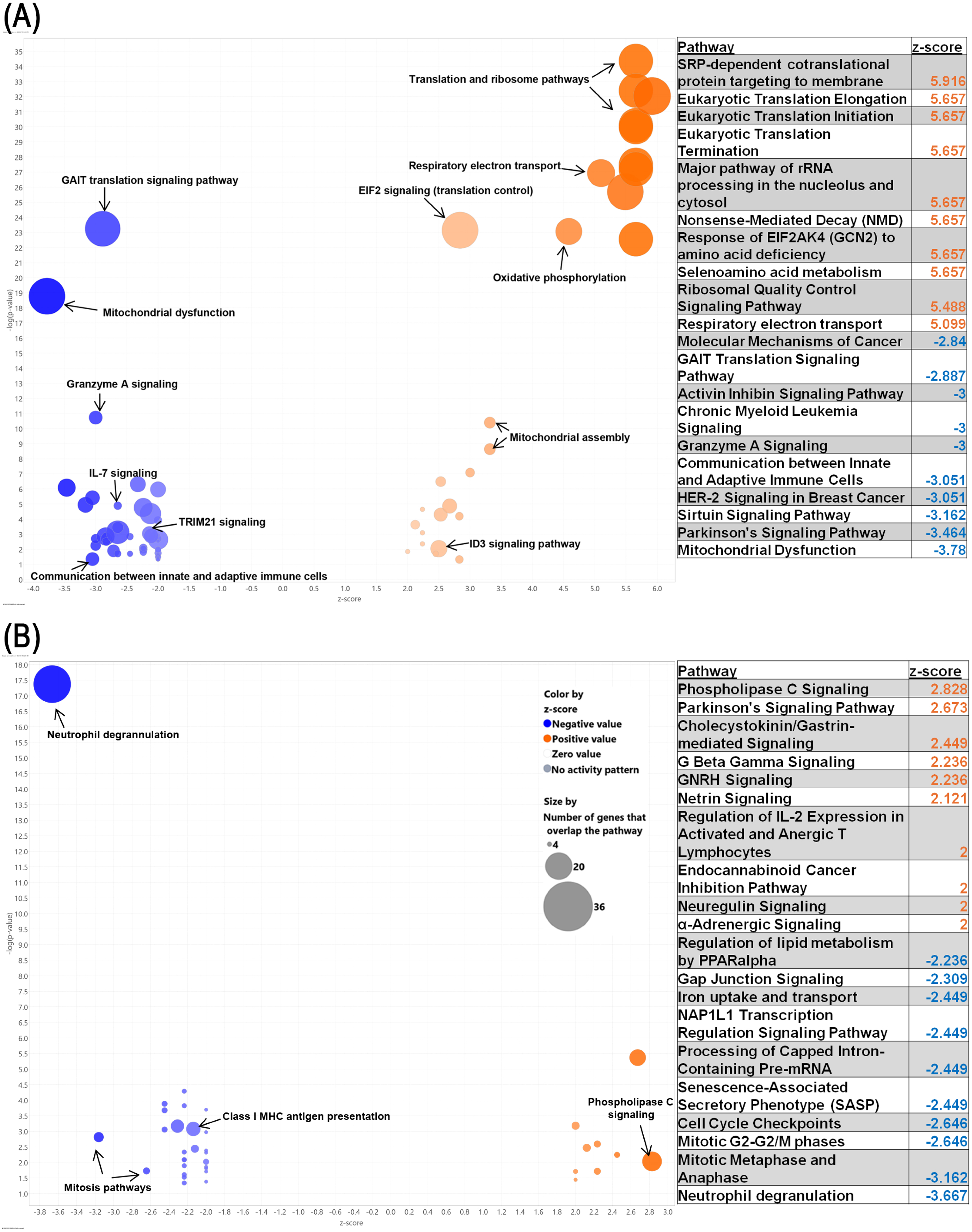
Age and Sex Pathway Analysis. **(A).** IPA analysis showed upregulation of pathways in the young mice compared to older mice. Pathways analysis of PMN-MDSCs are shown as a representative and showed upregulation of pathways regarding transcription/translation and mitochondrial assembly and function while downregulated pathways included mitochondrial dysfunction. Immune and inflammatory pathways displayed a more complex pattern with some upregulated pathways (i.e., neutrophil degranulation and ID3 signaling) and some downregulated (i.e., innate-adaptive immune communication, TRIM21 signaling, and IL-7) in the young adult septic mice compared to older. **(B)** IPA analysis for female sepsis vs male sepsis was performed and showed a mixture of up and down regulation. In PMN-MDSCs, as shown in this figure, more of the pathways were downregulated in the MDSCs from septic females compared to septic males. These included pathways associated with immune function and the cell cycle as shown in this volcano plot of pathways from PMN-MDSCs.

Sex-stratified analyses within the septic cohort revealed relatively few DEGs from all the MDSC subtypes between female and male adult mice (**Table 1**). However, IPA revealed distinct mixed pathway regulation patterns (**Fig. 2B**). Septic adult female mice demonstrated downregulation of neutrophil degranulation in all MDSC subsets. Notably, all pathways associated with protein synthesis, post-translational modifications, and cell cycle progression were downregulated in septic adult female MDSCs relative to septic adult males. Mitochondrial metabolic pathways were not prominently altered. These findings suggest sex-specific differences in immune signaling and protein homeostasis during sepsis.

In terms of combined age and sex effects, M-MDSCs had the most DEGs between all cohorts, demonstrating differences between older female and older male mice in addition to differences between young and older male mice (**Table 1**). Looking beyond individual genes, IPA illustrated that multiple immune and inflammatory response pathways were downregulated in septic older adult females compared to septic older adult male mice with neutrophil degranulation as the most significantly downregulated pathway in all MDSC subsets (**Fig. S2A**).

Similar to the sexual dimorphism revealed by comparing septic older adult females and males, there were mostly downregulated pathways when comparing young septic adult females and males (**Fig. S2B**). These included multiple pathways associated with protein synthesis/modifications and the cell cycle, immune and inflammatory responses, and mitochondrial and metabolism pathways.

In septic adult young female compared to septic older adult female mice there was a mixture of up- and down-regulation of IPA pathways (**Fig. S2C**). Pathways associated with mitochondria and metabolism were both up and downregulated; particularly, mitochondrial function was upregulated while mitochondrial dysfunction was downregulated in young adult septic female derived MDSCs (compared to older septic adult female MDSCs). All pathways associated with translation and ribosome function were upregulated in MDSCs of septic young adult females compared to those of septic older adult females.

There were more significant upregulated pathways in the MDSCs of the septic young adult male mice compared to septic adult older male mice in E- and M-MDSCs whereas there was a mixture of up and downregulated pathways in PMN-MDSCs (**Fig. S2D**). Pathways associated with protein synthesis, DNA repair, mitochondrial assembly, and the cell cycle were upregulated in the septic young males compared to septic older males whereas mitochondrial dysfunction related pathways were downregulated.

#### 3.2.3 Age- and sex-dependent divergence in myeloid differentiation trajectories revealed by pseudotime analysis

Pseudotime trajectory analysis revealed marked differences in age-dependent differences in myeloid cell expansion following sepsis (**Fig. 3**). Examining all splenic myeloid subsets revealed complex interactions between cell lineages regarding age, sex, and sepsis status (**Fig. 3**). As expected, naïve older adult mice exhibited enhanced baseline MDSC numbers compared to naïve young adults (e.g. inflammaging)^38–40^. After sepsis young adult females showed the most distinct differentiation pattern compared to septic young adult males, septic older adult males, and septic older adult females. Specifically, young adult female myeloid cells were predicted to differentiate more towards terminal cells (e.g. PMNs, monocytes), while the other three cohorts were predicated to dedifferentiation towards ‘stemness’ (e.g. MDSCs).

**Figure 3.**
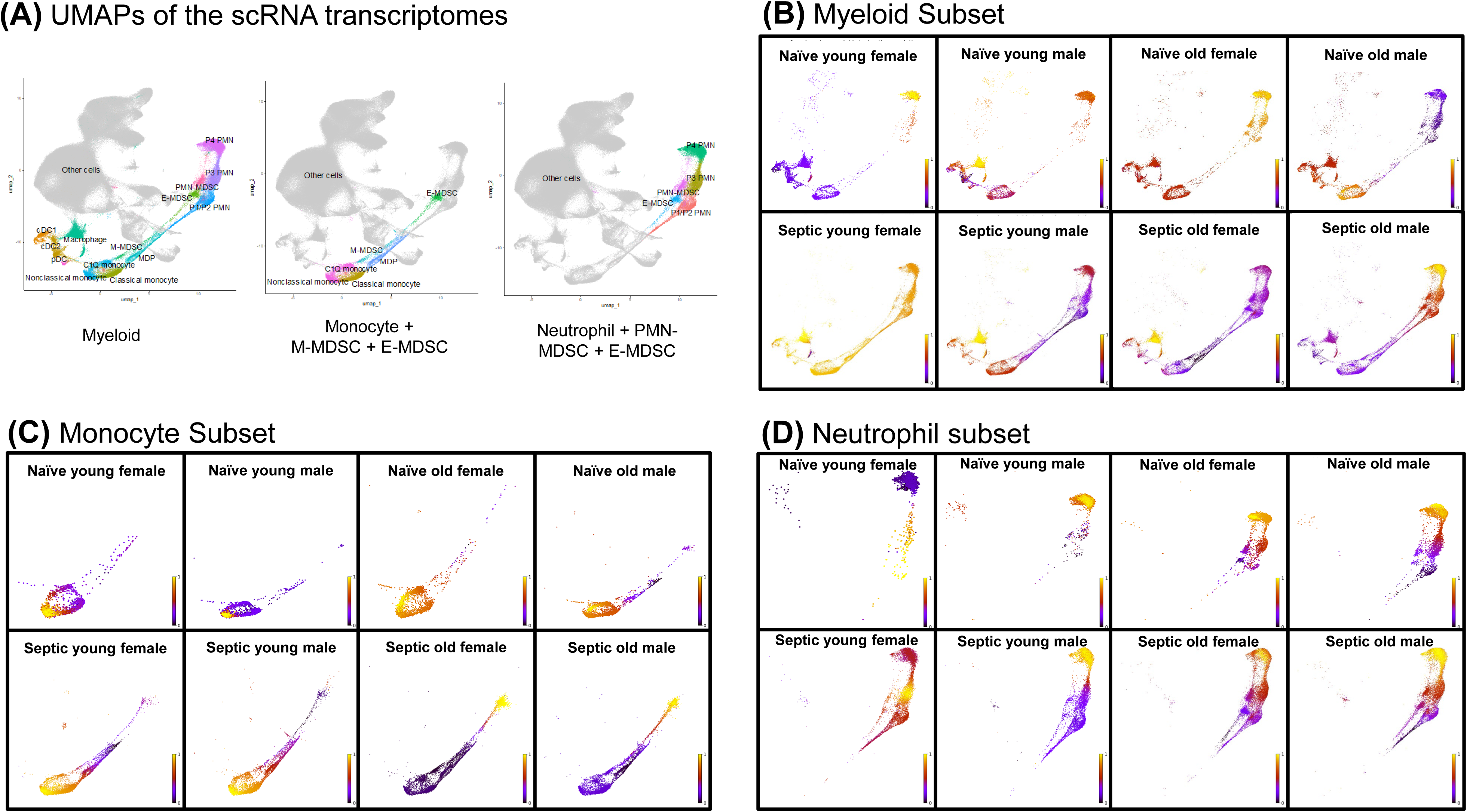
Pseudotime analysis. Pseudotime analysis (RNA velocity) was performed in the different cohorts. In the pseudotime analysis, purple indicates the “ancestor” and the more yellow indicates the cells are more differentiated. **(A)** UMAP embeddings of the scRNA transcriptomes of the myeloid compartment and the monocyte and neutrophil subsets. **(B)** The myeloid subset showed differences between naïve (top row) and septic (bottom row) as well as among the cohorts. **(C)** The monocyte subset showed opposite directionality between the young septic and old septic mice. **(D)** The young septic female mice differentiated to the P3 PMNs in the neutrophil subset, while the other septic mice differentiated to P4 PMNs.

To more precisely delineate predicted lineage differentiation, we performed pseudotime analysis in the two major developmental pathways: monocytic and granulocytic (**Figure 3A**). Within the monocytic subset, young adult septic and older adult septic mice displayed opposite directionality of differentiation, with the young adult mice favoring differentiation towards monocytes while the older adult mice differentiating towards E-MDSCs (**Fig. 3C**). Within the granulocytic lineage, our results illustrated notable divergences between male and female septic mice (**Fig. 3D**). Specifically, septic male mice exhibited a pronounced shift toward terminally differentiated neutrophils (P4), suggestive of accelerated or enhanced granulopoietic maturation in response to sepsis. In contrast, septic female mice maintained a broader distribution across intermediate states, including E-MDSCs and immature PMNs, indicative of a potentially more regulated or restrained granulocytic differentiation program. In sum, both age and sex, separately or combined, are predicted to alter host myelopoiesis after severe infection in unique ways, with young adult females globally demonstrating the most unique patterns.

### 3.3 Age- and sex-specific MDSC–lymphocyte communication networks in sepsis revealed by CellChat

Given the well-established immunosuppressive functions of MDSCs, and how age and sex are known to influence immune cell composition, signaling networks, and functional outputs during systemic inflammation, we next sought to examine how sepsis alters communication between MDSCs and T lymphocytes using Cellchat^41^.

Both age and sex predicted significantly different communication pathways between MDSCs as well as different lymphocyte clusters (**Fig. 4**). For example, PMN-MDSCs from older adult males showed significantly greater predicted interactions with CD8^+^ T cells than those derived from young adult males (*p*=0.03, **Fig. 4A**) after sepsis, which validates previous findings^19^. PMN-MDSCs from septic older adult females, however, showed significant greater interactions with CD4^+^CD25^+^ regulatory T cells (Treg) (*p* =0.017) and CD4^+^ T cells (*p* =0.03) compared to those from septic young adult females (**Fig. 4B**). M-MDSC interactions with T lymphocytes were different when stratified by sex: septic older adult male M-MDSCs had significantly greater interactions with CD8^+^T cells (*p*=0.014) versus septic older adult females (**Fig. 4C**). Additionally, septic young adult female E-MDSCs had greater interaction with B cells compared to septic older adult females (*p*=0.04) (**Fig. 4D**). Thus, MDSC lymphocyte interactions are influenced by age and sex.

**Figure 4.**
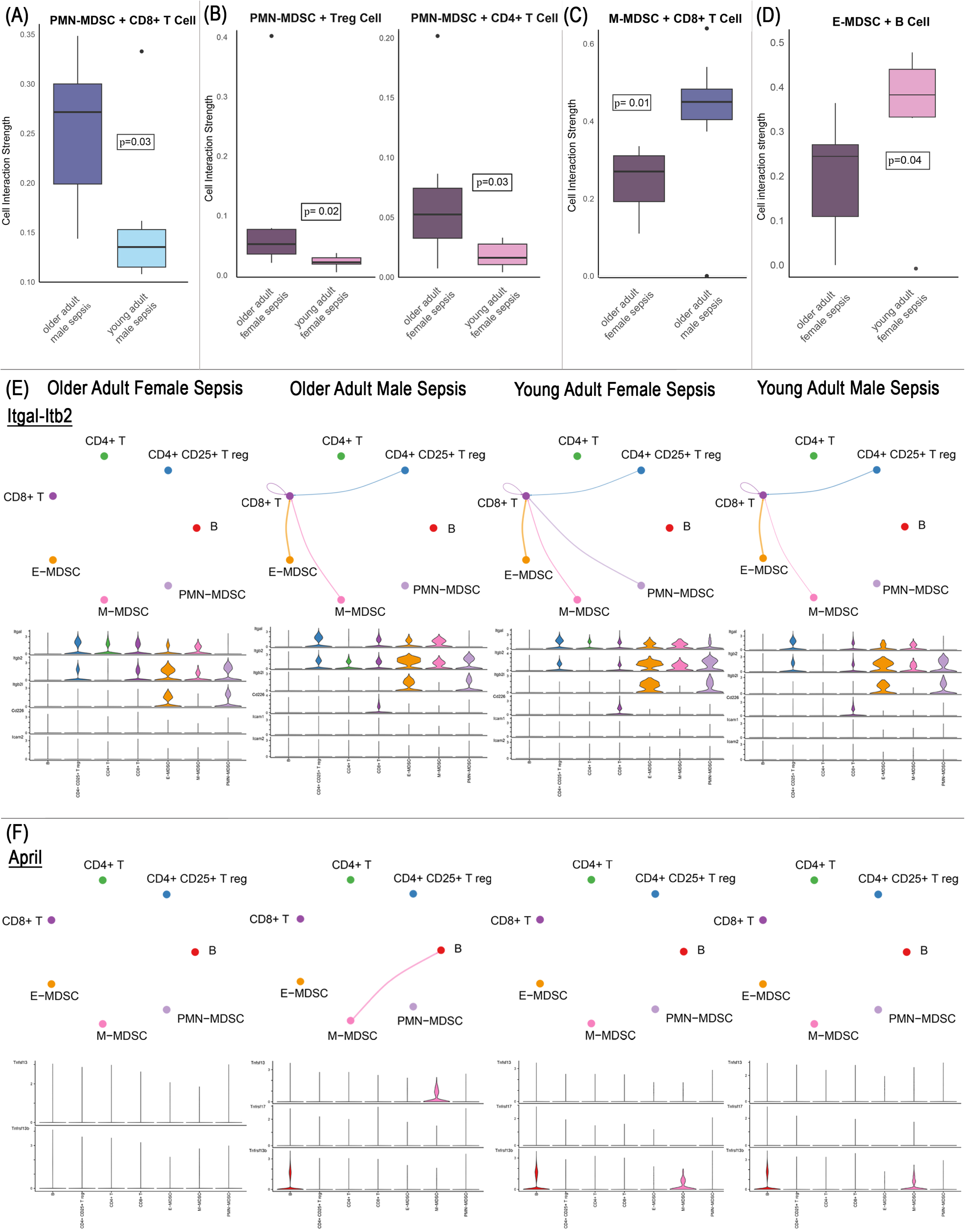
Predicted cell-cell interactions using Cellchat. **(A).** PMN-MDSCs from septic older adult male mice had greater predicted interactions with CD8+ T lymphocytes compared to septic young adult male mice. **(B)** PMN-MDSCs from septic older adult female mice had greater interactions with Treg lymphocytes and CD4+ T lymphocytes compared to septic young adult females. **(C)** M-MDSCs from septic older adult males had greater interactions with CD8+T lymphocytes than septic older adult females. **(D)** E-MDSCs from septic young adult females had greater intercations with B lymphocytes than septic older adult females. **(E)** MDSCs from young females had different interactions along the Itgal-Itb2 pathway compared to the other cohorts. There was higher gene expression of the ligand receptors in the young adult female PMN-MDSCs compared to the other cohorts as well as shown in the violin plots. **(F)** Septic older males, young females, and young males had unique communication patterns in the galectin pathway. **G)** Septic older males had a unique interaction in the April pathway and expressed both the ligand and receptor as seen in the violin plot.

These unique cohort responses are distinct regarding predicted pathways of communication in these cells as well. **Fig. 4E-F** highlights specific ligand-receptor pathways that are differentially enriched across cohorts as well as the strength of their receptor and ligand gene expression. Young adult female mice showed distinct communication between MDSCs and lymphocytes in multiple pathways including the Itgal-Itb2, Annexin, and Granulin pathways (**Fig. 4E, Fig. S3**). While septic older adult males had unique communication in the April pathway (**Fig. 4F**). This is corroborated by higher gene expression of the respective ligand-receptor pairs for these pathways compared to the other cohorts (violin plots, **Fig. 4E-F, Fig. S3**). In the galectin pathway, MDSCs from septic young females, septic young males, and septic older males had unique communication patterns (**Fig S3C**). Interestingly, CD8^+^ T lymphocytes from septic young females had similar communication patterns as older male mice (**Fig. S3C**).

### 3.4 Age and sex-dependent metabolic function of CD11b^+^Gr1^+^ splenocytes in septic mice

As previously stated, leukocyte mitochondria are central for appropriate cell function via multiple pathways^42^. Given the significant transcriptional upregulation of metabolic pathways observed in septic MDSCs compared to MDSCs derived from naïve mice, particularly those related to oxidative phosphorylation, glycolysis, and iron metabolism (**Fig. 1D**), we sought to functionally validate these findings by directly assessing mitochondrial and glycolytic activity in splenic CD11b^+^Gr1^+^ cells using extracellular flux analysis (**Fig. 5**). In disease states, CD11b^+^Gr1^+^ murine cells are comprised primarily of MDSCs as pathological signals induce their expansion and functional reprogramming^43^, while in naive mice these cells tend to be normal myeloid precursors. Also, our previous work has revealed that only CD11b^+^Gr1^+^ cells derived from older adult mice can immunosuppress lymphocytes seven days after CLP+DCS, meeting the criteria for MDSCs^19^.

**Figure 5.**
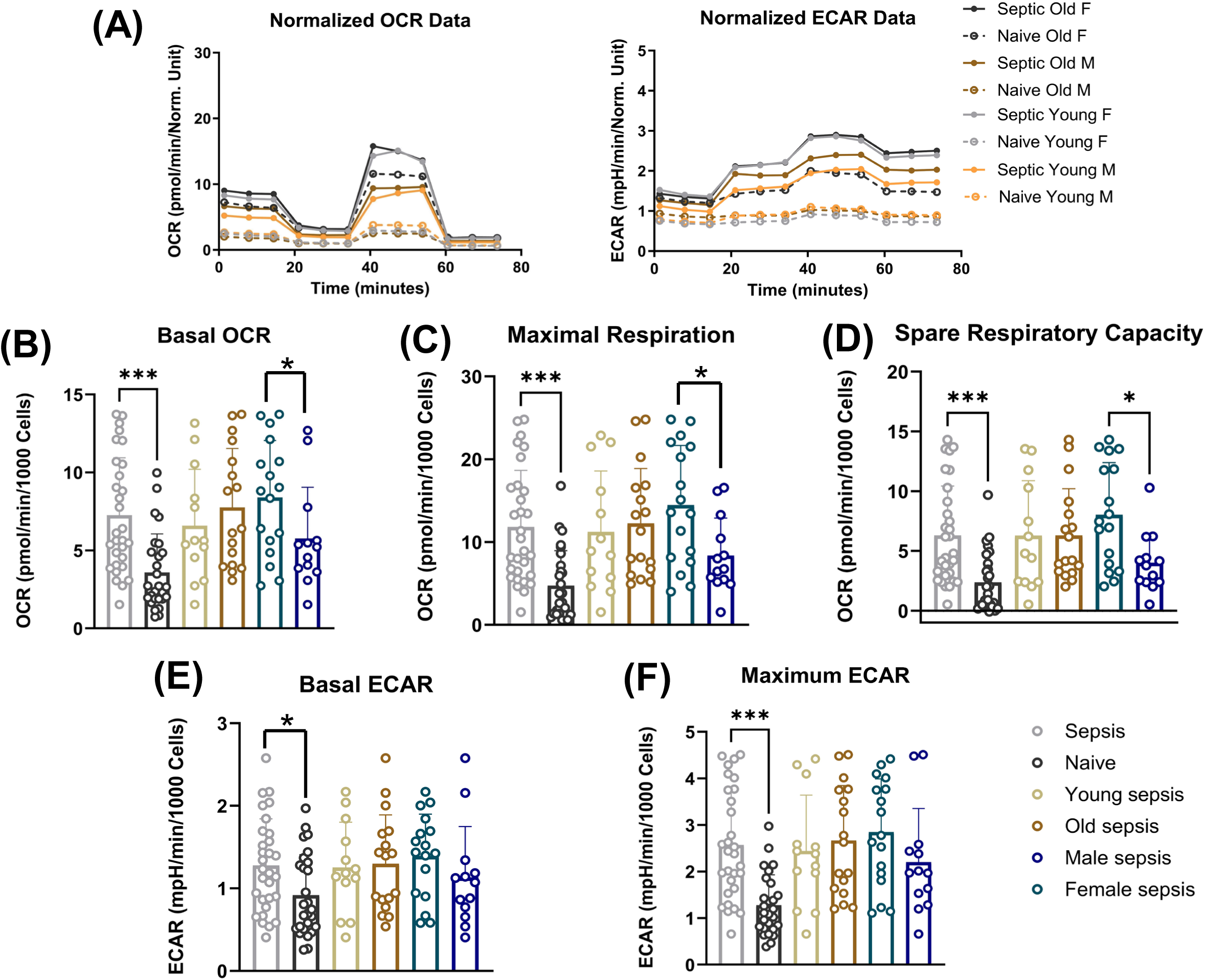
Mitochondrial metabolism of CD11b+Gr1+ cells. After sepsis, splenic CD11b+Gr1+ cells demonstrated increases in both oxygen consumption and glycolysis. **(A)** The average Seahorse traces by cohort for oxygen consumption (OCR) and extracellular acidification rate (ECAR) are shown. CD11b+Gr1+ cells from septic young females had pronounced increases in their metabolic activity compared to their naïve counterparts with increases in basal OCR, maximal respiration, spare respiratory capacity (SRC), and maximal ECAR in sepsis compared to naïve. Older males had increases in sepsis as well in these cells with increased basal OCR, maximal respiration, and maximum ECAR in sepsis compared to naïve. CD11b+Gr1+ cells from septic older females had greater maximal respiration and SRC compared to the older male mice. When examining trends from the larger groups, CD11b+Gr1+ cells from all septic mice had significantly greater basal OCR **(B)**, maximal respiration **(C)**, spare respiratory capacity (SRC) **(D)**, basal ECAR **(E)**, and maximum ECAR **(F)** than naïve mice. CD11b+Gr1+ cells from septic female mice had greater basal OCR **(B)**, maximal respiration **(C)**, and SRC **(D)** than those from septic male mice.

CD11b^+^Gr1^+^ cells from septic mice exhibited significantly elevated oxygen consumption compared to naïve. Basal OCR, maximal respiration, and spare respiratory capacity (SRC) were all increased. There were also significant increase in glycolysis with increased basal ECAR and maximal ECAR in CD11b^+^Gr1^+^ cells from septic mice compared to those from naïve mice (**Fig. 5, Table S5**)^44^. In summary, septic CD11b^+^Gr1^+^ cells exhibit a metabolically activated phenotype characterized by enhanced mitochondrial respiration and glycolytic capacity, consistent with a high-energy, immunoregulatory state that distinguishes them from their naïve counterparts.

Interestingly, although only splenic CD11b^+^Gr1^+^ cells from older adult septic mice previously met the functional definition of MDSCs, there were no significant differences in metabolism detected by the Seahorse assay when septic animals were stratified by age. However, CD11b^+^Gr1^+^ cells from septic adult female mice exhibited significantly higher mitochondrial respiratory activity compared to their male counterparts (**Fig. 5B-D, Table S5**). Notably, there were no significant sex-based differences in glycolytic activity, as measured by ECAR (**Fig. 5E, F**).

When examining combined effects, CD11b^+^Gr1^+^ cells from young adult female mice exhibited a pronounced increase in metabolic activity in response to sepsis, including elevated basal OCR, maximal respiration, and maximal ECAR, compared to naïve. This metabolic activation in young adult females paralleled the response seen in older adult mice, likely accounting for the lack of observed age-related differences in the pooled comparisons (**Fig. 5A, Table S5**). Additionally, septic older adult female CD11b^+^Gr1^+^ cells had significantly greater maximal respiration and SRC compared to septic older adult males (**Fig. 5A, Table S5**).

### 3.5 Predicted cohort specific drug target expression with splenic leukocytes based on transcriptomic expressions revealed by drug2cell

One key aspect to precision medicine is determining which patient cohorts may respond to certain drugs better than others. To continue to highlight this importance of precision therapy based on age and sex, we utilized drug2cell to predict drug targets based on the expression of drug targets in the single cell transcriptomics of all splenic white blood cells^45^. Multiple drugs were predicted to target septic splenic total leukocytes in different cohorts: young=26, older=123, female=32, male=42.

Given the use of steroids in sepsis remains controversial^46,47^ and is being studied for precision medicine use^48,49^, we specifically focused on differences identified with dexamethasone as the predicted target in our cohorts. In sepsis, males showed a higher targeting effect of dexamethasone compared to naïve (log2FC=0.23, *p* =0), while females trended to have a low targeting effect compared to naïve (log2FC=−0.301, *p* =0.07). This would imply that there may be sexual dimorphism in the host response to steroids after sepsis. There was decreased potential therapeutic cellular interaction of dexamethasone in the older adult septic mice (log2FC=−0.016, *p* =0.001) and no difference in the young adult septic mice. Again, this would imply that host responsiveness to steroids after sepsis could also be age dependent.

We next selected to evaluate Tretinoin, which is used to treat acute promyelocytic leukemia in patients. Moreover vitamin A derivatives have been found to decrease MDSC proliferation in HBV infected mice^50^ as well as inhibit some murine cancer progression via hastening MDSC differentiation to more mature immune cells^51,52^. In fact, all-trans-retinoic acid is being investigated as an anti-inflammatory agent and a mechanism to inhibit the expansion of MDSCs in sepsis^53–55^. Using drug2cell analysis of all the splenic leukocytes it was revealed that older adult septic mice were found to have increased targeting effect of tretinoin compared to their naïve state (log2FC=0.373, *p* <0.001) while there is no significant difference in the young cohorts. This would imply that MDSC therapeutics may be most effective in specific groups and should be investigated accordingly.

## Discussion

In over three decades, little to no progress have been made towards identifying an effective immunotherapeutic for sepsis^56^. Additionally, there has been recent justified criticism of preclinical-rodent sepsis models^57,58^. Historically, investigators have conducted research that was “cleaner” regarding delivering a standard effect on the host (e.g. LPS, no antibiotics, no fluid resuscitation) as well as standardizing the type of host utilized so that other factors (e.g. sex, age) did not confound the biological and pathological response^58–60^. Although the data generated was outstanding work regarding inflammation biology, individuals would often try to translate this research to humans without success^61^. More recently, medical scientists have embraced the concept of precision medicine, with the understanding that there is no “silver bullet” for sepsis^62^ and that any preclinical animal models need to not only embrace the complexity of sepsis, but mimic the current human environment in the surgical intensive care unit (e.g. fluid and antibiotics, stress, pain control, general anesthesia)^19,31,58,60^. Our work has attempted to address these issues by utilizing a pre-clinical murine surgical sepsis model that better imitates human severe infection^19^ with the specific intent of determining how age and sex can influence the myeloid response of the host. Using scRNAseq and metabolic profiling we demonstrated that both age and sex significantly influence the host immune response to severe abdominal infection. Notably, these factors critically affect the abundance, phenotype, and function of MDSCs, underscoring the necessity to consider biological context when evaluating immune responses and therapeutic strategies in sepsis^63^. **Figure 6** summarizes the key findings in this report emphasizing the effects of age and sex on the MDSC response to sepsis in the adult mouse.

**Figure 6.**
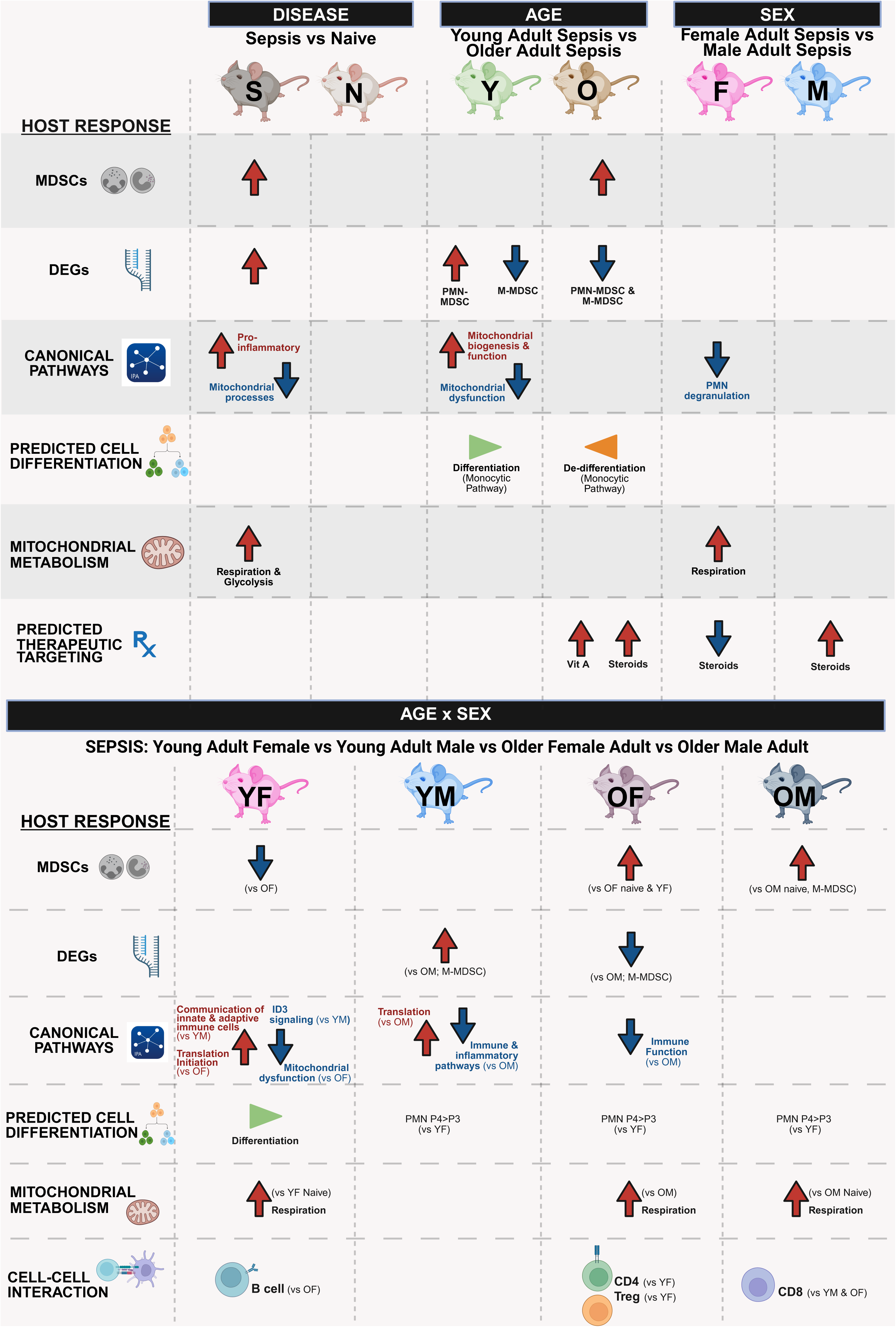
Summary of findings. Summary of main differences between sepsis vs naïve, septic older mice vs septic young mice, septic females vs septic males, and the cohorts. S, sepsis; N, naïve; Y, young; O, older; F, female; M, male; YF, young female; YM, young male; OF, older female; OM, older male. Created in https://BioRender.com

As post-sepsis in-hospital mortality has significantly decreased, we elected to use a model that induced sepsis (per Sepsis-3) but generated what has been defined as the Persistent Inflammation Immunosuppression Catabolism Syndrome (PICS) rather than acute mortality. Our work and others have demonstrated that MDSCs are associated with both post-sepsis PICS as well as poor outcomes in humans^1,3,4,10,29,64^. This work revealed that the MDSC response varies depending on the age and sex of the host. Importantly, this includes overall populations, transcriptomics, predicted differentiation and cell to cell interaction as well as mitochondrial function (**Fig. 6**).

Our work revealed that naïve older adult animals had enhanced baseline myelopoiesis compared to naïve young adults, which was expected given the known differences in cell proportions due to inflammaging and immunosenesence^19,65,66^. Additionally, there was a greater expansion of MDSCs in older adult mice after sepsis^19^, in agreement with previously published literature^19,38^. Surprisingly, we did not identify differences regarding MDSC prevalence between the sexes in both the septic and naïve animals.

When both age and sex were considered in septic animals regarding differential expression and pathway analysis, increased age (but not sex) displayed a relative “downregulation” of the MDSC transcriptome. This revealed an overall pattern of post-septic myelodyscrasia in the elderly. This indicates that restoring immune homeostasis to older adults will require a more nuanced intervention (e.g. not necessarily blocking inflammation) to improve outcomes across the spectrum of patients with sepsis and negative post-septic outcomes.

Differentiation within the myeloid compartment varied between both young and aged septic mice, as well as between sexes, with young adult female mice having the most pronounced differences (**Fig. 6**). RNA velocity of MDSCs in the context of sepsis remains largely unexplored but our findings suggest that the observed gene expression dynamics in these cells may underlie improved outcomes observed in young mice, particularly young females, potentially due to a more favorable differentiation trajectory that supports recovery.

Older adult mice had upregulated signaling pathways involved in communication between innate and adaptive immune cells. This was further supported by cell-cell communication analysis, which revealed increased interactions with CD8^+^, CD4^+^, and CD4^+^CD25^+^ regulatory T lymphocytes. Given that MDSCs contribute to immunosuppression by impairing T cell function and inducing T cell apoptosis, these heightened interactions may reflect or contribute to greater immune dysregulation in older mice^67^. While older adult mice showed increased interactions amongst various T lymphocyte populations, MDSCs from young adult females exhibited enhanced communication along specific pathways, particularly with CD8^+^ T cells via signaling with LFA-1, a key integrin involved in leukocyte adhesion and transmigration^68^. Females are known to mount more robust immune response across a range of diseases, including infections, often correlating with improved survival outcomes^69^. These findings underscore the complex role of MDSCs in sepsis, emphasizing that their function is not governed by a simple on/off switch in gene expression or cell-cell communications.

It has been previously published that MDSCs switch to glycolysis for energy production to meet their functional demands in the acidic environment in sepsis, as well as fatty acid oxidation, contributing to their immunosuppressive properties^70,71^. The fatty acid oxidation helps produce immunosuppressive cytokines and results in increased oxygen consumption^71^. In this study, MDSCs from septic female mice used oxygen at a greater rate and had a greater energy reserve (SRC) than those derived from male mice when stressed. Additionally, young female mice MDSCs had the greatest increases in both oxygen consumption and glycolysis compared to their naïve state. These findings suggest that female MDSCs may engage in a more robust mitochondrial metabolism following sepsis, potentially reflecting enhanced immunoregulatory or reparative functions. This sex-dependent metabolic reprogramming may be more functionally than transcriptionally regulated in this context given there were not significant mitochondrial transcriptomic changes between the sexes (as a whole). These findings suggest that the metabolic reprogramming of MDSCs in sepsis may be more influenced by sexual dimorphism than age.

The future of precision medicine lies not only in understanding the differences between cohorts but also targeting drugs to specific patients or cohorts of patients based off these differences. Utilizing drug2cell we demonstrated that there are predictions for drug targets regarding age and sex. Specifically, there were differences for dexamethasone regarding age and sex. Steroid use in sepsis remains controversial topic with mixed evidence regarding their effectiveness^72–74^. Although our work predicted therapeutics in murine total splenic leukocytes is not meant to affect current clinical practice, it does illustrate the importance of considering host factors when trying to conduct such translational work.

### Limitations

There were multiple limitations to this study. Firstly, we elected to not standardize the estrus cycle in the female mice and thus cannot comment on the specific contribution of estrogen or progesterone to the sepsis response. Sexual dimorphism arises not only from variations in circulating sex hormone levels but also from the differential expression of both hormone-dependent and hormone-independent genes, as well as from the host’s prolonged exposure to these mediators of gene expression^69^. Secondly, this study uses a preclinical murine model, so while it can be used as a basis to study sepsis in ways that are not feasible in humans, organoids, or by artificial intelligence, appropriate caution must be taken if trying to directly translate specific genes and proteins to humans. Thirdly, a notable limitation of this study lies in the use of the 10X Genomics scRNAseq platform, which, while enabling high-throughput single-cell transcriptomics, is known to capture only a subset of the transcriptome, particularly favoring highly expressed genes. This bias can limit detection of low-abundance transcripts and potentially affect the comprehensive characterization of rare cell populations. Future studies employing complementary approaches, such as CITE-seq or targeted sequencing, may provide more comprehensive insights into the functional states of MDSCs in sepsis. Fourthly, the results from the metabolic assays were obtained on total CD11b^+^Gr1^+^ cells compared to the rest of the analyses focusing on individual myeloid subsets. Fifthly, mice were euthanized at the same time-point and sepsis is well known to be a dynamic process after infection. Future studies will need to also consider such variables as time, type of infection and organ of infection, as these also influence the host leukocyte response and septic outcomes^75–78^. Finally, drug2cell has been validated in humans but it is unknown how this translates to murine cells, though we can assume that the genes evolved from a common ancestral gene so likely perform similar functions. However, the role of drug2cell in this work was to further highlight the importance of considering sex and age in sepsis research rather than to identify specific drugs for clinical trials.

Despite these limitations, altogether, we determined that the MDSCs in survivors of sepsis who develop a PICS phenotype are indeed unique based on host factors such as age and sex. Combined, our data illustrates that a precision medicine approach will be required to modulate MDSC activities in the septic host.

## Resource Availability

Requests for further information and resources should be directed to and will be fulfilled by the lead contact, Philip A. Efron, M.D.

## Materials availability

This study did not generate new unique reagents

## Data and Code availability

Data pending submission to GEO.

## Acknowledgments

This work was supported, in part, by the following National Institutes of Health grants:

NIH T32 GM-008721

RF1NS128626

NIA - R21AG087039

## Author Contributions

Conceptualization: PAE, GC, RM, LLM

Formal analysis: DW, HT, XY, LBN, CR, GC, PAE

Funding acquisition: PAE, GC, PC

Investigation: MLD, RU, DCN, LZS

Methodology: PAE, LLM

Project administration: PAE

Supervision: PAE, GC, LLM, RM

Visualization: CR, DW, HT, PAE

Writing – original draft: CR, DW, HT, MLD, DCN, LZS, GC, PAE

Writing – review & editing: CR, DW, HT, VEP, MHR, LBN, JCR, AC, SF, QV, MLD, DCN, LZS, LLP, SDL, GC, FX, SMW, MAB, TJL, LB, AMM, PC, MPK, CEM, LLM, RM, GC, PAE

## Methods

### Experimental Model

#### Mice

All studies were approved by the Institutional Animal Care and Use Committee at the University of Florida. C57BL/6J (B6) mice were purchased from Jackson Laboratories (Bar Harbor, ME) and housed at the University of Florida’s (UF) Animal Care Services facility, where they received standard care: *ad libitum* feeding and 12-hour light-dark cycle schedule. Mice were housed together for no less than two weeks to allow their microbiome to equilibrate^79^.

#### Sepsis Induction

Naïve and septic, young adult (3-4 months) and older adult (18-24 months) female and male B6 mice were used for all studies (**Suppl. Table 1**). Sepsis was induced using the cecal ligation and puncture with daily chronic stress (CLP+DCS) model^19,28^. Seven days post-CLP+DCS (CLP+DCS 7d) animals were euthanized to harvest splenocytes^19,31^.

## Methods Details

### Single-cell RNA sequencing (scRNA-seq) analysis

#### Single cell collection

Splenocytes were obtained and counts were quantified using a Cellometer™ Auto 2000 Cell Viability Counter (Nexcelom Bioscience, Lawrence, MA). One million cells with a minimum of 85% cell viability were obtained for single cell RNA sequencing (scRNAseq) using 10X Genomics chemistry (Chromium X instrument, Pleasonton, CA). Libraries were prepared with the 10X’s Chromium Next GEM Single Cell 5’ v2 kit, profiling 10,000 cells per sample. To ensure quality during library preparation, RNA integrity and double-stranded cDNA insert size were assessed using the TapeStation 4200 (Agilent, Santa Clara, CA) and Qubit 4 (Invitrogen, Carlsbad, CA). Final concentration of the library pool was measured with the NEBNext Library Quant Kit (New England Biolabs, Ipswich, MA). Libraries were sequenced at a minimum depth of 20,000 read pairs per cell on an Illumina NovaSeq X plus Platform (Illumina, San Diego, CA).

#### Data pre-processing

Cell Ranger Single-Cell Software v 7.0.1 (10X Genomics) was used to obtain FASTQ files from the Illumina base call files. The reads were aligned to the mm10 mouse transcriptome reference using “intronic reads included” mode, with an expected cell count of 10,000. Transcript expression was quantified at the single-cell level, generating gene-cell count matrices.

#### Quality control, integration and clustering

The Cell Ranger-generated count matrices were imported into R (v4.3.2) and analyzed using the Seurat (v5.0.1) package^80^. Quality control (QC) steps were applied to remove low-quality cells (cells expressing > 10% mitochondrial transcripts or number of unique genes < 200) and doublets (number of unique genes > 6000). Gene expression data were log normalized and scaled. Principal Component Analysis (PCA) was performed based on the top 2,000 most variable genes. To account for batch effects across different samples, data integration was performed using the Harmony (v1.2.0) package^81^, using the top 50 principal components (PCs) to align datasets and reduce batch-specific variations. A shared nearest neighbor graph was constructed using 30 nearest neighbors based on the top 30 PCs. Further, cell clusters were detected based on the Louvain algorithm and were visualized in Uniform Manifold Approximation and Projection (UMAP) embeddings computed from the top 30 PCs.

#### Cell-type annotation

Cell types were manually annotated by assigning clusters to known cell types based on marker genes, following previous studies^82^. First, broad cell types were identified: T-cells, B-cells, plasma cells, dendritic cells, macrophages, monocytes, and neutrophils (PMNs). Within the myeloid cell compartment, re-clustering with adjusted resolution was performed to resolve final cell subsets.

### Statistical Analysis

#### Models for group comparisons

To evaluate the effects of disease status, age, and sex on gene expression and cell-type proportions and other features of interest, we employed a hierarchical modeling strategy using generalized linear models. This strategy allowed us to (1) assess the overall impact of sepsis compared to naïve while adjusting for host factors (age and sex); (2) isolate the individual contributions of age and sex within each disease state (sepsis and naïve). Specifically, for each feature (e.g., gene expression, cell proportion), the outcome *y_ij_* in a particular cell type *j* of subject *i* was modeled as

Level 1:

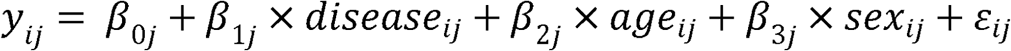

Level 2:

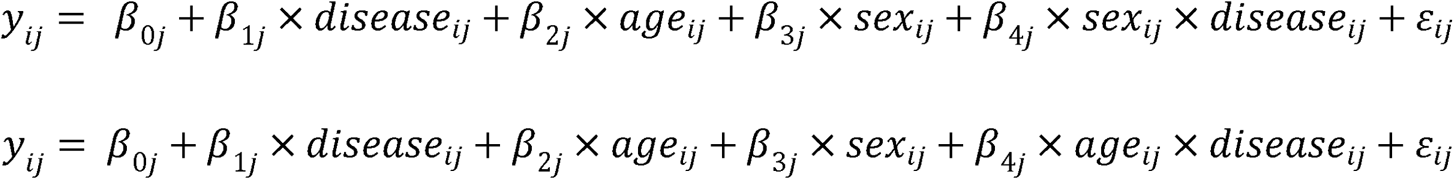

Level 3:

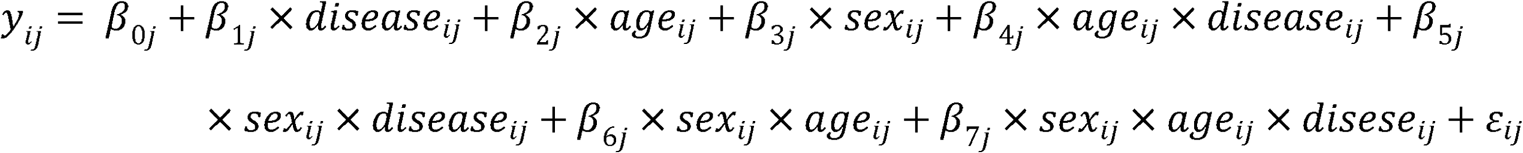

In each case, β denotes the mean changes in gene expression corresponding to disease status (sepsis vs naïve), age (older vs young), sex (female vs male) and their interactions. *ε_ij_* is the random error with variance *σ*^2^. The statistical analysis was performed using R (v4.3.2). The outcome *y_ij_* was transformed when appropriate (e.g., log transform for gene expression count). Multiple testing was addressed by applying the Benjamin-Hochberg correction and a false discovery rate (FDR) *q* < 0.05 was considered significant.

#### Cell-type proportion differential analysis

Cell-type proportions in each sample were calculated and compared using the three-level approach as shown in 2.4.1, for (1) sepsis vs. naïve, (2) older vs. young and female vs. male, and (3) more granular contrasts between specific age × sex × disease subgroups, e.g., young female sepsis vs. young female naïve.

#### Pseudobulk differential expression analysis

The *Aggregate Expression* function in Seurat was used to aggregate gene expression per cell type within each sample. R package edgeR (v4.0.16) was further used for differential expression analysis. The pseudobulk sequencing count data was modeled as described in *Models for group comparisons,* with the log link function applied.

#### Pathway enrichment analysis

Enriched canonical pathways were identified using Ingenuity Pathway Analysis (IPA QIAGEN Inc. Germantown, MD, https://digitalinsights.qiagen.com/IPA) using Fisher’s exact test and adjusted with the Benjamini-Hochberg correction on detected differentially expressed genes (DEGs). If fewer than 300 DEGs were identified, the top 300 genes with the lowest FDR were used for analysis to avoid potential bias from the size of the input. Significance was determined using the activation z-score (z-score ≥ 2 refers to predicted activation and z-score ≤ −2 refers to predicted inhibition for the canonical pathways identified).

#### Velocity analysis

To infer the dynamic transcriptional trajectories of MDSC subsets in sepsis, RNA velocity analysis was performed using Python 3.10.13 and scVelo (v0.3.1)^83^, based on the pre-mature (unspliced) and mature (spliced) transcript information, which were obtained by Velocyto (v0.17.17)^84^. Preprocessing included selection of the top 2,000 highly variable genes with expression in at least 20 cells, followed by computation of first- and second-order moments using the first 30 PCs and the 30 nearest neighbors. Pseudotime was visualized on UMAP embeddings derived from the Seurat pipeline.

#### Cell-cell communication analysis

Using the scRNA-seq data, we inferred the communication probabilities between cell types based on overexpressed ligand-receptor interactions by CellChat (v2.1.0)^85^. Cell-cell communication networks were constructed and visualized in circular network plots, after filtering out low-confidence interactions. Circular network plots illustrate each signaling pathway’s contribution, with edge thickness proportional to interaction strength. Using the three-level modeling approach described above, the differences in MDSC:T/B cell communication probabilities were compared among experimental groups.

#### Drug2cell of total splenic leukocytes to determine cohort differences

To identify potential therapeutic targets specific to different host cohorts, we used drug2cell (v0.1.1)^45^ to quantify relationships between drugs and drug targets. To adapt human-centric drug target data for analyzing mouse data, we performed orthology mapping to convert human gene targets to their mouse homologs using Ensembl BioMart database^86^. We curated the ChEMBL dataset with HUGO Gene Nomenclature Committee (HGNC) lists of classes of drug targets, such as ion channels, G-protein coupled receptors, kinases, and nuclear hormone receptors. Of note, total leukocytes were used as drug2cell is designed to score drug-target interactions across annotated cell populations and our intent was to illustrate cohort specific results rather than isolate a “drug that could cure MDSCs/sepsis in a mouse”^45^. Filtering was performed to focus on FDA-approved drugs with functional assays documenting bioactivity below drug target thresholds taken from the Illuminating the Druggable Genome (IDG) project^87^. Wilcoxon rank sum tests were used to compare drug-target scores between sepsis and naïve groups within each subgroup (e.g., older, young, male, female), with Benjamini-Hochberg corrected *p* values.

#### Metabolic and extracellular flux analyses (Seahorse XF Analysis)

Similar to our previous work, we used the EasySep™ Mouse MDSC (CD11b^+^Gr1^+^) Isolation Kit to obtain murine splenic MDSCs^19^. It should be noted that this technique can isolate a mixed myeloid population, particularly in naïve mice. The metabolic capacity of splenic CD11b^+^Gr1^+^ cells (2.5 x 10^5^ per well) were analyzed using a Seahorse XFe96 Extracellular Flux Analyzer (Agilent Technologies, Santa Clara, CA) by means of mito-stress test assays performed according to the manufacturer’s protocol. Oxygen consumption rate (OCR) and extracellular acidification rate (ECAR) values were averaged between technical replicates and normalized by adding Hoechst 33342 dye (Cell Signaling Technology, Danvers, MA) for fluorescent cell counting with a Cytation 1 microplate reader (Agilent Technologies, Santa Clara, CA).

