## Supplemental material for "Age- and Sex- Driven Transcriptional and Metabolic Diversity in Myeloid-Derived Suppressor Cells After Mouse Sepsis"

**Supplemental Table 1** Number of mice analyzed per cohort.

| Group | N of samples |
|---|---|
| Young female naïve | 6 |
| Young female sepsis | 7 |
| Young male naïve | 8 |
| Young male sepsis | 7 |
| Old female naïve | 6 |
| Old female sepsis | 7 |
| Old male naïve | 8 |
| Old male sepsis | 8 |
| Total | N = 57 |

**Supplemental Table 2** Percentage of MDSC subtypes per cohort relative to all the cells isolated.

| *Cell Type* | **Young female naïve** | **Young female sepsis** | **Young male naïve** | **Young male sepsis** | **Old female naïve** | **Old female sepsis** | **Old male naïve** | **Old male sepsis** |
|---|---|---|---|---|---|---|---|---|
| **E-MDSC** | 0.00% | 0.17% | 0.01% | 0.26% | 0.10% | 1.42% | 0.03% | 1.06% |
| **M-MDSC** | 0.01% | 0.18% | 0.01% | 0.33% | 0.04% | 0.69% | 0.01% | 0.57% |
| **PMN-MDSC** | 0.04% | 0.42% | 0.06% | 0.59% | 0.39% | 2.82% | 0.15% | 1.61% |

**Supplemental Table 3** Proportion of the MDSCs in the disease status, age, and sex comparisons.

| | | **E-MDSC** | **M-MDSC** | **PMN-MDSC** |
|---|---|---|---|---|
| **Sepsis vs naïve** | *z* score | 2.82 | 4.49 | 3 |
| | Adjusted *p* value | **0.01* | **0.0001* | **0.008* |
| **Young sepsis vs old sepsis** | *z* score | -2.84 | -2.51 | -2.88 |
| | Adjusted *p* value | **0.01* | **0.023* | **0.009* |
| **Female sepsis vs male sepsis** | *z* score | -0.12 | 0.39 | -0.71 |
| | Adjusted *p* value | 0.897 | 0.696 | 0.483 |

**Supplemental Table 4** Cell proportion analysis by cohort and MDSC subtype

| Comparison | | E-MDSC | M-MDSC | PMN-MDSC |
|---|---|---|---|---|
| Young female sepsis vs young female naive | z-score | 0.370 | 0.828 | 0.560 |
| | adj p-value | 0.996 | 0.680 | 0.837 |
| Young male sepsis vs young male naive | z-score | 0.633 | 2.007 | 0.845 |
| | adj p-value | 0.996 | 0.112 | 0.746 |
| Old female sepsis vs old female naive | z-score | 2.330 | *3.084* | *2.690* |
| | adj p-value | 0.072 | **0.013* | **0.027* |
| Old male sepsis vs old male naive | z-score | 2.255 | *3.018* | 1.954 |
| | adj p-value | 0.072 | **0.013* | 0.152 |
| Young female sepsis vs old female sepsis | z-score | -2.272 | *-2.509* | *-2.762* |
| | adj p-value | 0.072 | **0.040* | **0.027* |
| Young male sepsis vs old male sepsis | z-score | -1.643 | -0.997 | -1.290 |
| | adj p-value | 0.251 | 0.597 | 0.422 |
| Young female sepsis vs young male sepsis | z-score | -0.232 | -1.046 | -0.259 |
| | adj p-value | 0.996 | 0.597 | 0.854 |
| Old female sepsis vs old male sepsis | z-score | 0.464 | 0.514 | 1.295 |
| | adj p-value | 0.996 | 0.900 | 0.422 |

**Supplemental Table 5** P values for the mitochondrial metabolism analysis of CD11b+Gr1+ cells. Red indicates that there was an increase in metabolism in the comparison (example sepsis had increased basal OCR compared to naïve) and blue indicates decreased metabolism.

| | **Basal OCR** | **Proton leak** | **Maximal respiration** | **Spare respiratory capacity** | **Basal ECAR** | **Maximal ECAR** |
|---|---|---|---|---|---|---|
| **Sepsis vs naïve** | <0.001* | 0.0004* | <0.001* | <0.001* | 0.047* | <0.001* |
| **F v m sepsis** | 0.03* | 0.017* | 0.004* | 0.001* | 0.297 | 0.108 |
| **Y vs o sepsis** | 0.326 | 0.401 | 0.683 | 0.901 | 0.857 | 0.562 |
| **YF vs YM sepsis** | 0.127 | 0.128 | 0.068 | 0.068 | 0.466 | 0.195 |
| **OF vs OM sepsis** | 0.174 | 0.075 | 0.028* | 0.009* | 0.825 | 0.410 |
| **YF vs OF sepsis** | 0.664 | 0.607 | 0.822 | 0.956 | 0.873 | 0.911 |
| **YM vs OM sepsis** | 0.463 | 0.695 | 0.860 | 0.852 | 0.825 | 0.637 |
| **YF sepsis vs naïve** | 0.002* | 0.004* | 0.0002* | 0.0002* | 0.095 | 0.002* |
| **YM sepsis vs naïve** | 0.174 | 0.108 | 0.112 | 0.126 | 0.556 | 0.129 |
| **OF sepsis vs naïve** | 0.214 | 0.453 | 0.067 | 0.095 | 0.825 | 0.108 |
| **OM sepsis vs naive** | 0.01* | 0.04* | 0.035* | 0.126 | 0.556 | 0.041* |

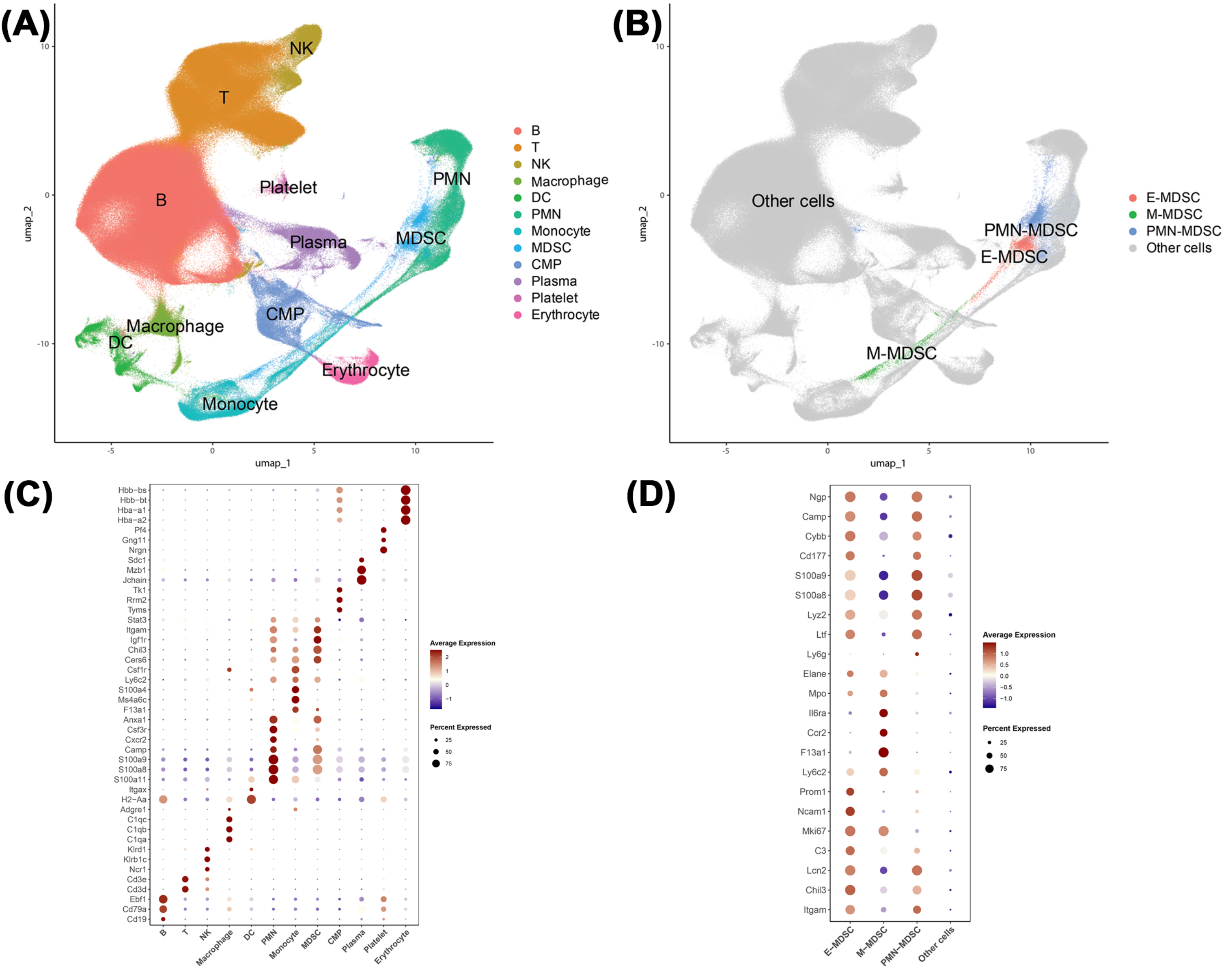


**Supplemental Figure 1** scRNA seq was performed and cell types visualized via UMAP **(A)** and annotated using known maker genes **(B)** starting with broad cell types. MDSCs were then classified as either early (E-), monocytic (M-), or granulocytic (PMN-) MDSCs **(C & D)**. Natural killer cells (NK), Dendritic cells (DC), Polymorphonuclear cells (PMN), Myeloid-derived suppressor cells (MDSC), Common myeloid progenitors (CMP)

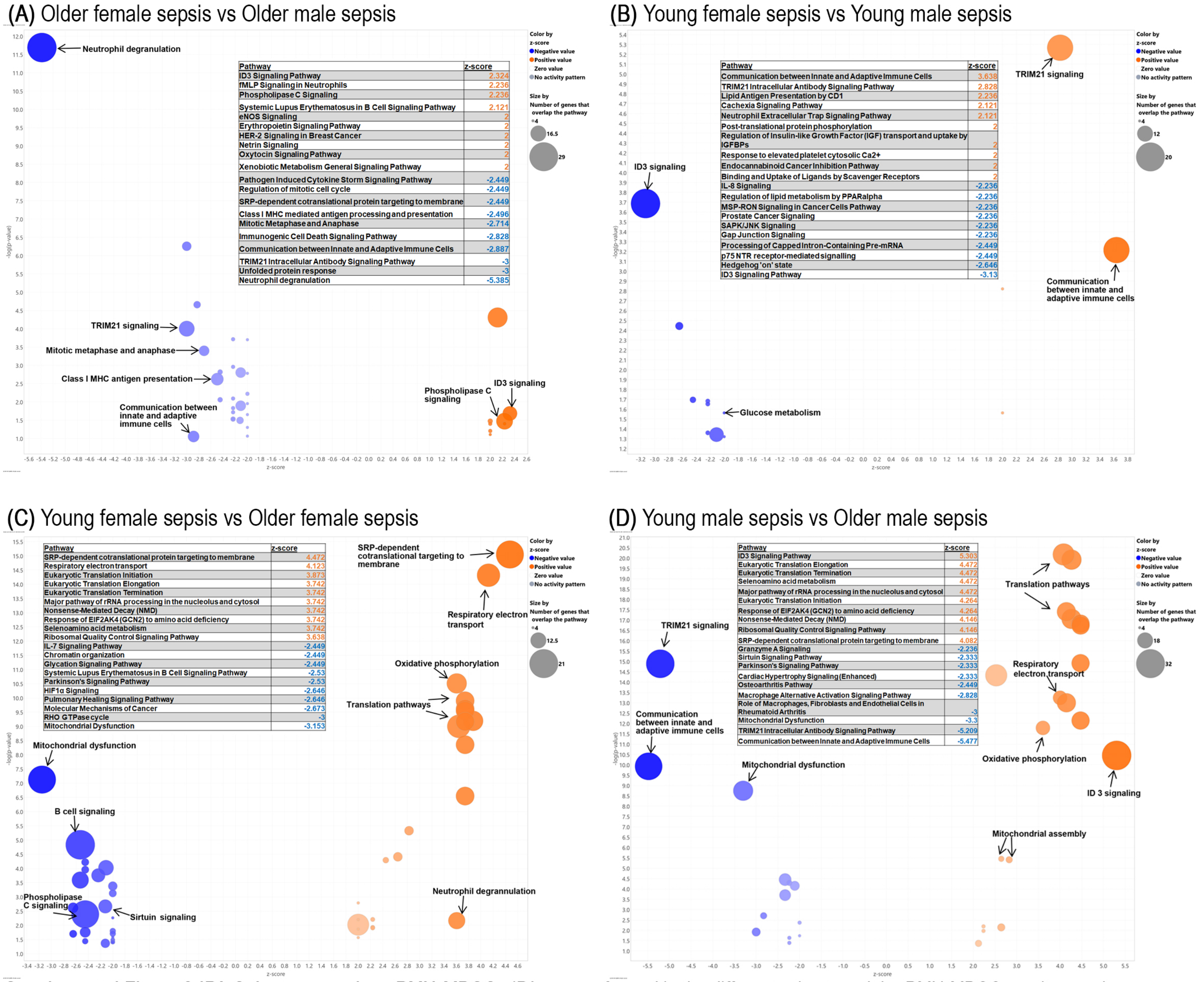


**Supplemental Figure 3 IPA Cohort comparison PMN-MDSCs** IPA was performed in the different cohorts and the PMN-MDSC results are shown as a representation of all the MDSCs. **(A)** More pathways were downregulated in the MDSCs from older adult females compared to septic older adult males. Neutrophil degranulation was the most significantly downregulated pathway in all MDSC subsets. There were multiple other pathways associated with immune and inflammatory responses that were downregulated in the older adult females compared to males like communication between innate and adaptive immune cells and MHC antigen presentation. Multiple pathways associated with protein synthesis/modification and the cell cycle were also downregulated in the older females for example, cell cycle checkpoints and post-translational protein modifications. Few pathways were significantly upregulated in the septic older female mice compared to septic older adult males. **(B)** There were mostly downregulated pathways when comparing MDSCs from young adult septic females compared to young adult septic males. These included multiple pathways associated with protein synthesis/ modifications and the cell cycle (translation pathways, DNA synthesis, nonsense mediated decay), immune and inflammatory response (MHC antigen presentation and IL-8 signaling), and mitochondrial and metabolism pathways (glucose metabolism and mitochondrial protein degradation). The upregulated pathways in septic young females compared to septic young males were mostly related to immune and inflammatory responses (e.g communication between innate and adaptive immune cells). **(C)** In MDSCs from septic adult young female compared to septic older adult female mice there was a mixture of up- and down-regulation of IPA pathways. Some pathways associated with mitochondria and metabolism were upregulated (oxidative phosphorylation, mitochondrial assembly) while mitochondrial dysfunction, granzyme A signaling, and sirtuin signaling were among those associated with metabolism that were downregulated in young septic females compared to older septic adult females. All pathways associated with translation and ribosome function (e.g ribosome quality control and translation initiation) were upregulated in septic young females compared to septic older females. Most pathways associated with immune and inflammatory processes were downregulated in the septic adult young females including IL-7 and T cell signaling and communication between innate and adaptive immune cells while neutrophil degranulation and ID3 signaling were upregulated in young adult females. **(D)** There were more significant upregulated pathways in the MDSCs from septic young adult male mice compared to septic adult older male mice in E- and M-MDSCs whereas there was a mixture of up and downregulated pathways in PMN-MDSCs. All pathways associated with protein synthesis, DNA repair, and the cell cycle were upregulated in the septic young adult male compared to septic older adult male MDSCs. Pathways associated with mitochondrial assembly and function were upregulated in the septic young male mice MDSCs and mitochondrial dysfunction related pathways were downregulated. Pathways associated with immune and inflammatory responses like communication between innate and adaptive immune cells were downregulated in septic young male MDSCs whereas ID3 signaling was upregulated in E- and PMN-MDSCs. Additionally related to immune responses, other interleukin and S100 family signaling were downregulated in PMN-MDSCs in septic young adult male mice.

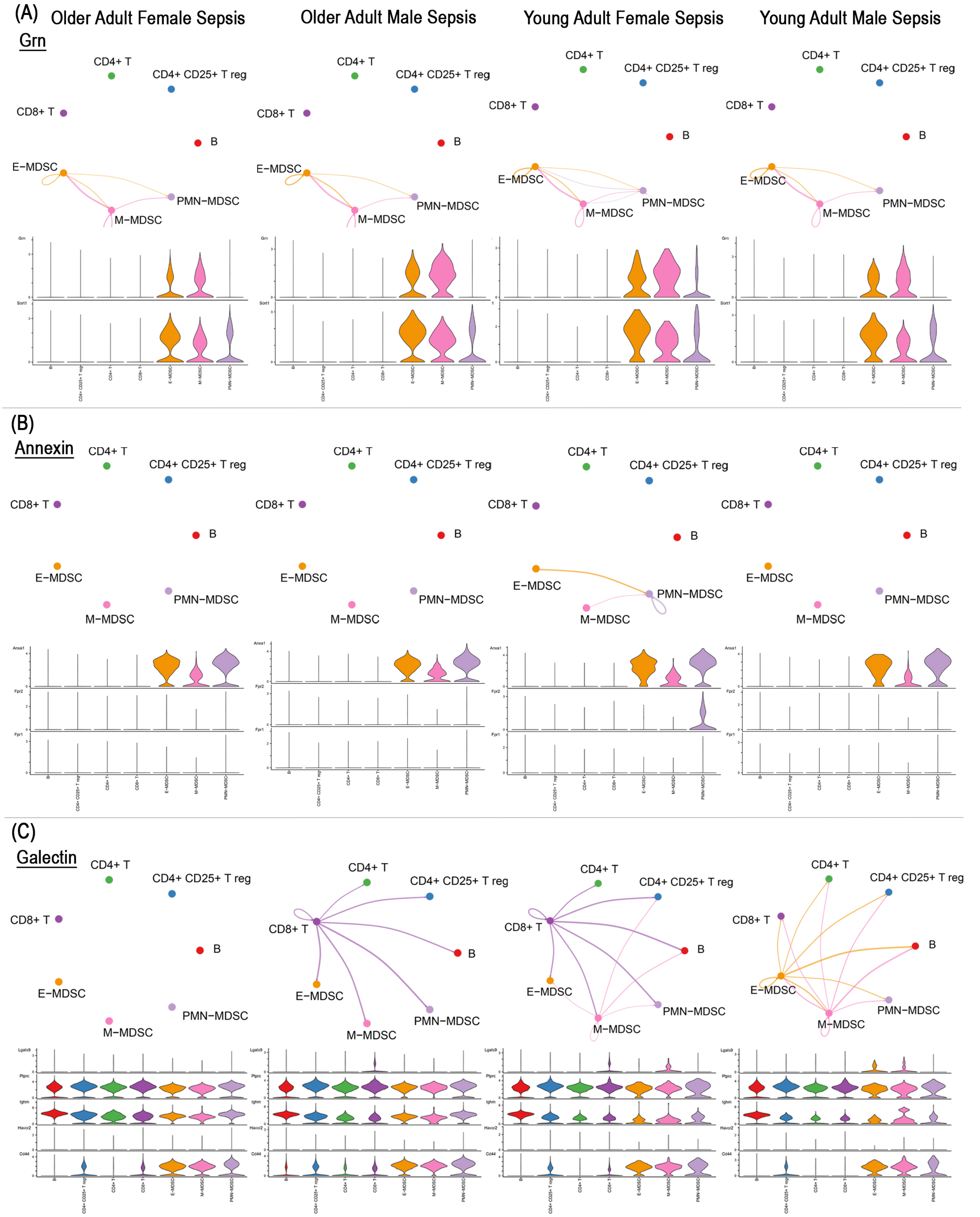


**Supplemental Figure 3 Further Cell Communication Interactions via Cellchat** MDSCs from septic young females had further unique interactions in the Grn **(A)** and Annexin **(B)** pathways. **(C)** There were unique interactions in young females (though similar to older males), and young males in sepsis in the Galectin pathway.
